# Recently evolved human-specific methylated regions are enriched in schizophrenia signals

**DOI:** 10.1101/113175

**Authors:** Niladri Banerjee, Tatiana Polushina, Francesco Bettella, Sudheer Giddaluru, Vidar M. Steen, Ole A. Andreassen, Stephanie Le Hellard

**Author notes:** **Correspondence to**: Prof. Stéphanie Le Hellard, Department of Clinical Medicine, Laboratory Building, Haukeland University Hospital, N-5021 Bergen, Norway. **Email**. **Authors’ email addresses** Niladri Banerjee.

## Abstract

**Background:** One explanation for the persistence of schizophrenia despite the reduced fertility of patients is that it is a by-product of recent human evolution. This hypothesis is supported by evidence suggesting that recently-evolved genomic regions in humans are involved in the genetic risk for schizophrenia. Using summary statistics from genome-wide association studies (GWAS) of schizophrenia and 11 other phenotypes, we tested for enrichment of association with GWAS traits in regions that have undergone methylation changes in the human lineage compared to Neanderthals and Denisovans, i.e. human-specific differentially methylated regions (DMRs). We used analytical tools that evaluate polygenic enrichment of a subset of genomic variants against all variants.

**Results:** Schizophrenia was the only trait in which DMR SNPs showed clear enrichment of association that passed the genome-wide significance threshold. The enrichment was not observed for Neanderthal or Denisovan DMRs. The enrichment seen in human DMRs is comparable to that for genomic regions tagged by Neanderthal Selective Sweep markers, and stronger than that for Human Accelerated Regions. The enrichment survives multiple testing performed through permutation (n=10,000) and bootstrapping (n=5,000) in INRICH (*p*<0.01). Some enrichment of association with height was observed at the gene level.

**Conclusions:** Regions where DNA methylation modifications have changed during recent human evolution show enrichment of association with schizophrenia and possibly with height. Our study further supports the hypothesis that genetic variants conferring risk of schizophrenia co-occur in genomic regions that have changed as the human species evolved. Since methylation is an epigenetic mark, potentially mediated by environmental changes, our results also suggest that interaction with the environment might have contributed to that association.

## Background

Schizophrenia is a psychiatric disorder that has been reported throughout human history, possibly as far back as 5000 years [1, 2]. Family, twin and adoption studies estimate that schizophrenia has a high heritability of 60-90% [3–6]. Today, schizophrenia is estimated to have a prevalence of 1%. It is associated with reduced fertility and increased mortality [7–11], and its persistence despite this heavy burden is paradoxical. Power et al [11] leveraged Swedish registry data to demonstrate the reduced fecundity of patients with schizophrenia, despite the novel finding that sisters of individuals with schizophrenia had higher fitness than controls. They henceforth suggested hitherto unknown mechanisms for persistence of the disease. One explanation for this persistence is that evolution has indirectly selected the disease instead of eliminating it - the disease may co-segregate with creativity and intellectual prowess, providing selective advantages to the kin of affected individuals [9, 12]. Crow first argued that language and psychosis may have common origins, which could explain the persistence of schizophrenia in human populations [12, 13]. This evolutionary hypothesis of the origins of schizophrenia can now be tested, thanks to the identification of genetic factors implicated in schizophrenia [14–16] and the availability of datasets that reflect recent genomic evolution in humans [17–19].

Large genome-wide association studies (GWAS) have identified thousands of variants that are associated with schizophrenia [14–16] but our mechanistic understanding of the candidate variants is poor. One approach to investigating the function of schizophrenia-associated variants is comparative genomics, which investigates the evolutionarily relevance of certain genomic regions [20]. This field has introduced new datasets to test disease origins in humans, including Human Accelerated Regions (HARs) and Neanderthal Selective Sweep (NSS) scores [18, 19]. HARs are genomic regions that are highly conserved in non-human species, but have undergone rapid sequence change in the human lineage [20–24]. Xu et al [18] showed that genes near HARs are enriched for association with schizophrenia. Neanderthals were hominids that co-existed and even bred with modern humans [25, 26]. Comparison of Neanderthal and human genome sequences [27, 28] has revealed genomic regions that have experienced a selective sweep in modern humans, presumably following a favorable mutation [28]. Negative NSS scores can be used to pinpoint mutations (usually single nucleotide changes) that were positively selected in humans as they diverged from Neanderthals. Srinivasan et al [19] found that genomic regions tagged by negative NSS scores show enrichment of association with schizophrenia.

Using specific interpretation of genome sequencing in two recently extinct hominids, Neanderthals and Denisovans, Gokhman et al [29] mapped genome-wide methylation levels (i.e. the methylome) and compared them to modern humans. While 99% of the methylation maps were identical in the three hominids, nearly 2000 differentially methylated regions (DMRs) were identified, which give the first clues about the role of epigenomic evolution in generating anthropometric differences between modern humans and their ancient cousins [29]. These DMRs provide a dataset of evolutionary annotations complementary to pre-existing datasets. Unlike HARs and NSS scores, which are based on DNA sequence changes, DMRs provide information on the evolution of epigenomes. Since epigenomes can act as an interface with the environment [30, 31], these datasets provide the opportunity to investigate environmentally driven evolutionary changes. Keeping in mind the evolutionary hypothesis for schizophrenia proposed by Crow, we thus examined if these evolutionary DMRs are enriched for association with schizophrenia. We also examined a range of human traits to compare the possible enrichment in other traits. Using previously published methodologies [19, 32, 33] and publicly available GWAS datasets we systematically analyzed twelve diverse phenotypes to investigate the potential role of regions susceptible to epigenetic variation in the emergence of specific traits in the human lineage.

## Results

### SNPs in human-specific DMRs are enriched for association with schizophrenia

The genomic locations of human-specific DMRs were obtained from data published by Gokhman et al [29] (see Methods for full details). GWAS summary statistics for 12 common traits were obtained from published datasets: schizophrenia [14], bipolar disorder (BPD) [34], attention deficit hyperactivity disorder (ADHD) [35], rheumatoid arthritis [36], high density lipoprotein [37], low density lipoprotein [37], triglycerides [37], total cholesterol [37], systolic blood pressure [38], diastolic blood pressure [38], body mass index [39], and height [40]. The GWAS datasets are summarized in Additional File 1, Table S1. For each trait, we generated a list of single nucleotide polymorphisms (SNPs) within DMRs (positional annotation) and a list of SNPs in linkage disequilibrium (LD-based annotation) with markers within DMRs (Additional File 1, Table S1).

We used quantile-quantile (QQ) plots as described by Schork et al [32] to test whether the DMR SNPs are enriched for association with the GWAS trait compared to the complete set of SNPs (see Methods for additional details). In such plots the baseline is the null line of no difference between expected distribution of *p*-values and observed *p*-values. Deviation of the observed data distributions from the expected data distribution indicates the presence of true associations. When the *p*-values for a set of selected markers show greater leftwards deflection, they are enriched for association compared to the overall GWAS set. For the schizophrenia GWAS, enrichment was observed both for SNPs in LD with markers in DMRs (Figure 1; Additional File 1, Figure S1) and for SNPs located within DMRs (Figure 2). Although there was a slight leftward deflection in the higher *p*-values (smaller negative log_10_ of *p*-values) in some other traits (e.g. height; Figure 1; Additional File 1, Figure S1), the observed enrichment only crosses the genome-wide significance level of 5×10^−8^ for the schizophrenia SNPs. The enrichment of disease-associated markers in DMRs is thus specific to schizophrenia and is independent of LD.

**Figure 1:**
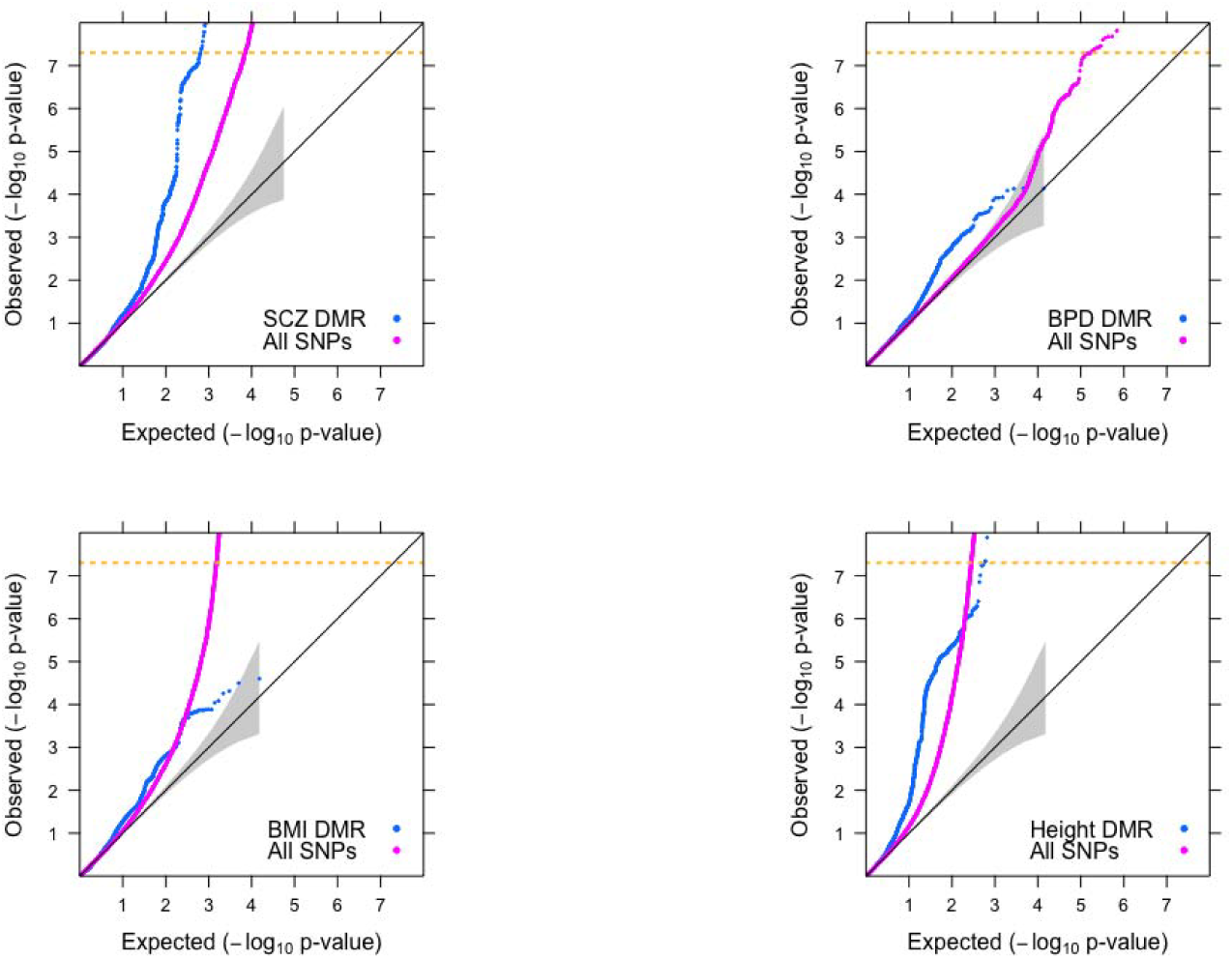
Enrichment of DMR SNPs across SCZ, BPD, BMI and Height. Quantile-Quantile (QQ) plots of GWAS SNPs for Schizophrenia (SCZ) with the extended MHC region masked (chr6: 25-35Mb), Bipolar Disorder (BPD), Body Mass Index (BMI) and Height. The X-axis shows expected -log_10_ *p*-values under the null hypothesis. The Y-axis shows actual observed -log_10_ *p*-values. The values for all GWAS SNPs are plotted in pink while the values for SNPs in linkage disequilibrium (LD) with DMRs are plotted in blue. Leftwards deflections from the null line (grey diagonal line) indicate enrichment of true signals - the greater the leftward deflection, the stronger the enrichment. Genomic correction was performed on all SNPs with global lambda.

**Figure 2:**
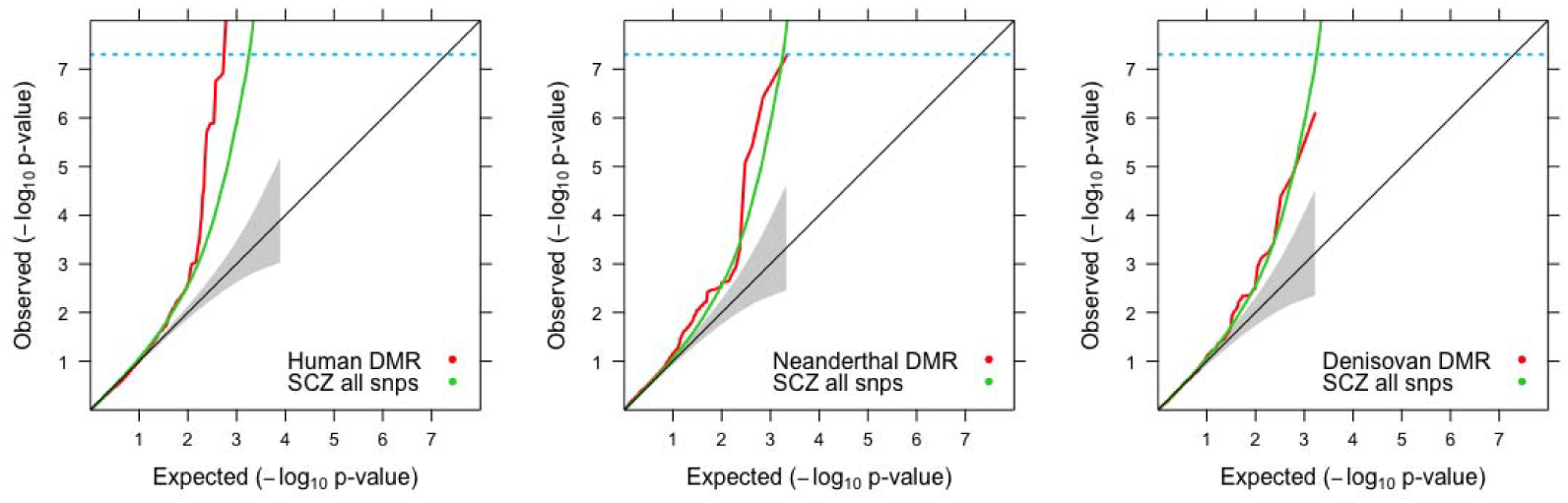
Comparison of enrichment of association with schizophrenia for SNPs within Human, Neanderthal and Denisovan DMRs. The figure shows QQ plots for all schizophrenia (SCZ) GWAS SNPs in green while SNPs within the species-specific DMRs are plotted in red. The location of the MHC is unknown in Neanderthal and Denisovan genomes, so the MHC region was not masked in the human genome.

### Human-specific DMR enrichment in schizophrenia is independent of the MHC region, other genomic annotations and total markers genotyped

The Major histocompatibility complex (MHC) region harbors several significant schizophrenia markers and could potentially bias our results because of long-range LD. The QQ plots show that the enrichment remains when the MHC is excluded (Figure 1) or included (Figure 2).

The schizophrenia GWAS had the highest density of markers genotyped (~9.4 million) and thus had the most SNPs in DMR regions (Additional File 1, Table S1), which could artificially inflate the enrichment. We normalized the total number of DMR SNPs with the total number of SNPs genotyped in each GWAS and found that the proportion of SNPs in DMRs is nearly identical for all traits (Additional File 1, Figure S3). To further eliminate the possibility that the enrichment is due to variation in the number of markers analyzed, we extracted ~2.4 million SNPs that were common across the twelve GWAS. Although not as strong as with the full set, the deflection observed for the schizophrenia GWAS remains higher than any other trait (Additional File 1, Figure S1), indicating the presence of significant disease markers in DMRs. These validations point to a true enrichment of association of the DMR SNPs with schizophrenia that is independent of the number of markers in a GWAS. It should be noted that we cannot rule out enrichment in the ADHD and BPD GWAS, because they are lacking in power (Additional File 1, Figure S1).

Additionally, we considered the distribution of schizophrenia-associated SNPs based on genomic annotations of 5’ untranslated regions (5’UTRs), Exons, Introns and 3’ untranslated regions (3’UTRs) [32]. Contrary to previously published findings [32], the enrichment was highest for intronic SNPs and lowest for 5’UTR SNPs (Additional File 1, Figure S4).

### Only human-specific DMRs are enriched for association with schizophrenia

Next, we used QQ plots to test whether markers located in the Neanderthal- and Denisovan-specific DMRs are enriched for association with schizophrenia. Coordinates for these DMRs were obtained from data published by Gokhman et al, 2014 [29] (see Methods for details). Since we do not know the precise coordinates of the MHC for Neanderthals and Denisovans, the analysis for human DMRs included the MHC region. No enrichment was observed for Neanderthal or Denisovan DMRs (Figure 2). It should be noted that this approach may not be appropriate for testing Neanderthal- and Denisovan-specific DMRs since (a) the schizophrenia GWAS was conducted in humans; (b) SNP and LD information is available only for humans; (c) the three hominids had variable number of DMRs, which affected the number of SNPs captured via positional annotation.

### Comparison of human DMRs with other evolutionary annotations

We compared the enrichment observed for the human DMRs with the enrichment previously reported for NSS markers and HARs [18, 19] (see Methods for details). We first compared the enrichment via QQ plots and find that the enrichment of human DMRs in schizophrenia is comparable to that observed for NSS markers and far greater than that observed for HARs (Figure 3).

**Figure 3:**
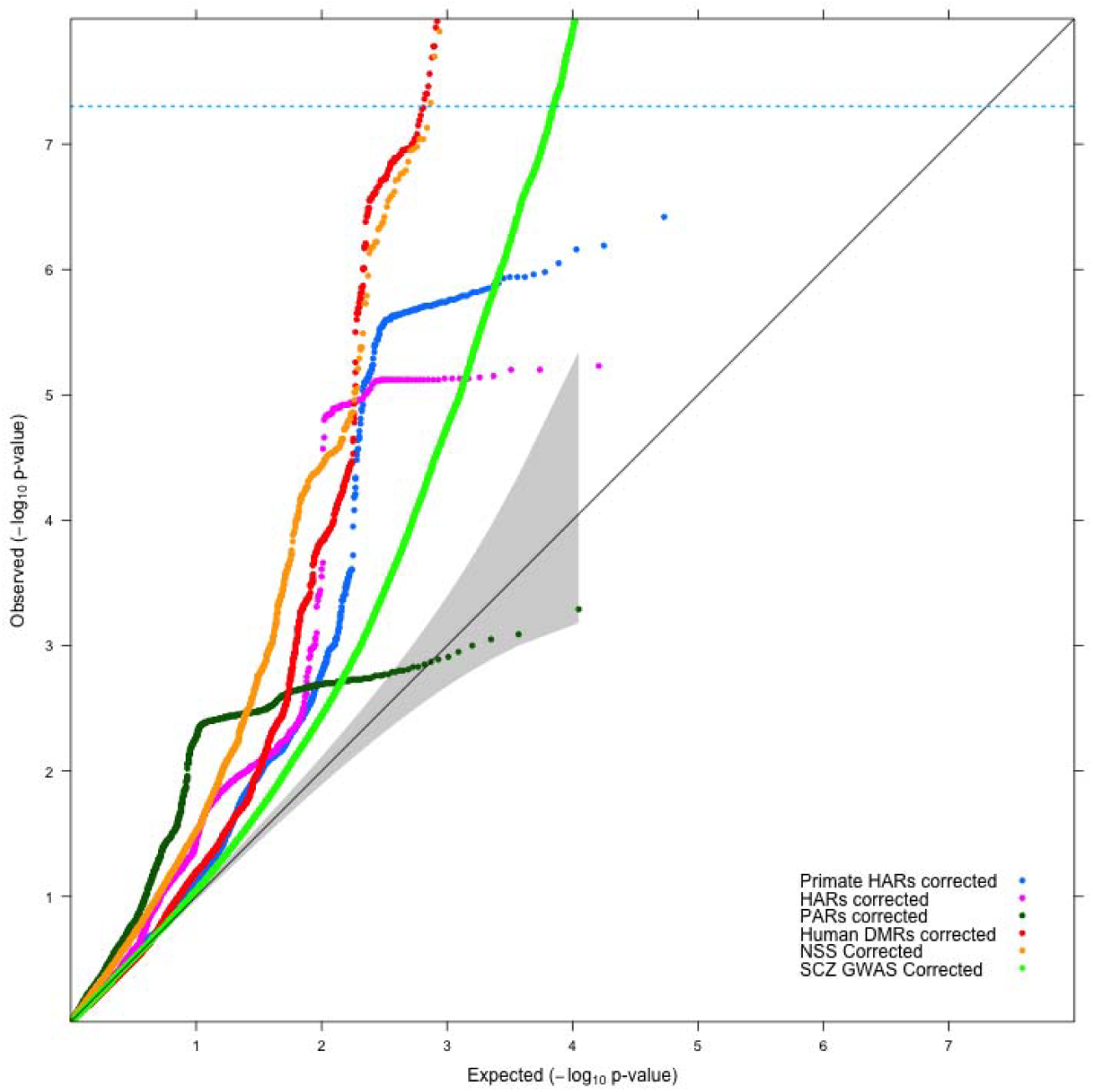
Comparison of enrichment of association with schizophrenia for SNPs in LD with various evolutionary annotations. QQ plots for association with schizophrenia (SCZ) of SNPs in different evolutionary datasets (DMRs - red, NSS - orange, Primate HARs (pHARs) - blue, HARs - magenta, PARs - dark green) versus schizophrenia GWAS with all SNPs (light green). SNPs are corrected for genomic inflation using global lambda.

In these analyses, it was important to check the extent of overlap of markers (SNPs) annotated to various genomic regions of DMRs, NSS markers and HARs. Reassuringly, the various evolutionary annotations do not share the same group of markers, indicating that we did not test the same regions or SNPs (Additional File 1, Figure S2). The overlap between NSS markers and DMR markers involved less than 0.5% of all NSS markers and less than 0.2% of all DMR markers (Additional File 1, Figure S2). The SNPs in the DMRs thus represent a different group of markers that have not been annotated or analyzed previously from an evolutionary standpoint (Additional File 2, Additional File 3).

### Statistically-significant enrichment exists for human DMRs

To determine the statistical significance of the DMR enrichment in schizophrenia, we utilized the INRICH software pipeline. INRICH is a pathway analysis tool for GWAS, designed for detecting enriched association signals of LD-independent genomic regions within biologically relevant gene sets (in our case genes which contain DMRs). It performs permutation and bootstrapping procedures to determine the significance of association of markers in LD intervals while maintaining the SNP density and gene density of the original intervals [33]. INRICH confirmed significant (*p*<0.05) enrichment of association for human DMRs with schizophrenia after correcting for multiple testing through bootstrapping at most *p*-value thresholds of LD intervals. Additionally, INRICH independently verified the previously reported enrichment of NSS markers with schizophrenia [19] (Figure 4). Furthermore, INRICH identified gene-level enrichment of association for DMRs with height (Additional File 1, Figure S5), while at the SNP level the enrichment in height was seen only for smaller effects, i.e. the enrichment did not remain below *p*<10^−8^.

**Figure 4:**
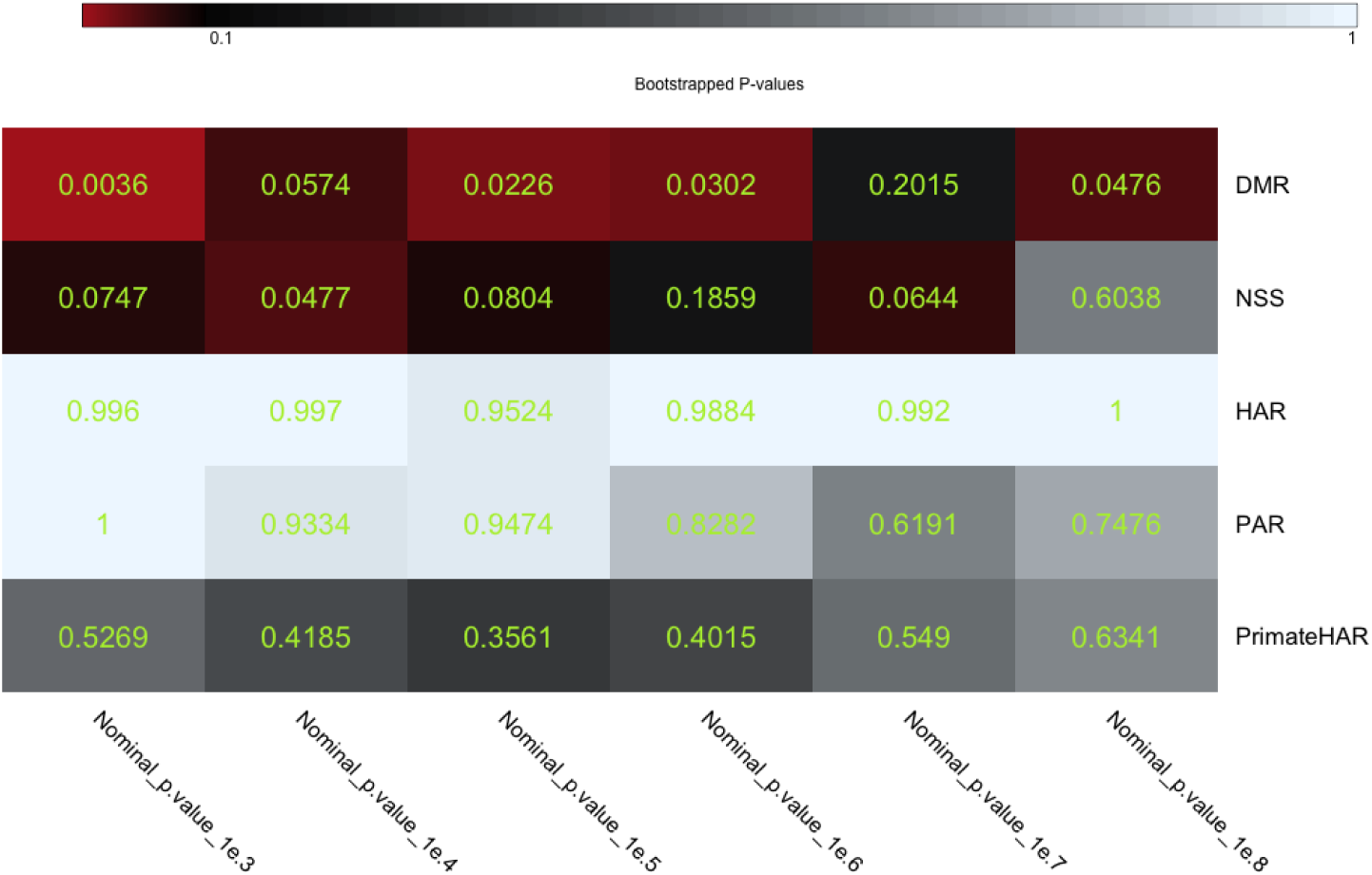
INRICH test for enrichment of association of DMR, NSS and Accelerated Region gene sets. Corrected *p*-values based on performing multiple testing with bootstrapping 5000 times, with *p* =0.1 as threshold. The various evolutionary annotations compared are: DMR, human-specific DMRs; NSS, Neanderthal Selective Sweep; HAR, mammalian conserved regions that are accelerated in humans; PAR, mammalian conserved regions that are accelerated in primates; and PrimateHAR (pHAR), primate-conserved regions that are accelerated in humans.

### Pathway analysis

We utilized Ingenuity Pathway Analysis (IPA) to analyze DMR SNPs that show enrichment of association with schizophrenia (for details of the genes analyzed, please refer to the ‘Pathway analysis’ section in the Methods). We found ‘CREB signaling in neurons’ and ‘Synaptic long term potentiation’ amongst the top canonical pathways when analyzing pathways overrepresented in nervous system. Additionally, under physiological systems, ‘Nervous system development and function’ is also enriched (Additional File 1, Table S2). We repeated the same analysis for NSS markers as they also show enrichment of association with schizophrenia. ‘CREB signaling in neurons’ was also amongst the top canonical pathways for enriched NSS markers (Additional File 1, Table S4). Additionally, we repeated the analyses with all organ systems and even then, ‘CREB signaling in neurons’ and ‘Synaptic long term potentiation’ emerged amongst the top canonical pathways for both enriched DMRs (Additional File 1, Table S3) and enriched NSS (Additional File 1, Table S5). This is an interesting result since there is very little marker overlap between the DMR and NSS SNPs (Additional File 1, Figure S2). Interestingly, genes containing enriched DMRs are also overrepresented in ‘Hair and skin development’ when considering all organ systems (Additional File 1, Table S3). This may suggest potential gene-by-environment interactions, modulated by methylation variation over human evolution (see Discussion below).

## Discussion

Our results suggest that SNPs in regions of the human genome that have undergone recent changes in DNA methylation status are enriched for association with schizophrenia, and to a lesser extent with height. Amongst all the traits analyzed, the enrichment observed in QQ plots was strongest for schizophrenia and passed the genome-wide significance threshold of 5×10^−8^ when the MHC was both excluded (Figure 1) and included (Figure 2). INRICH analysis confirms significant enrichment (*p*<0.01) in human DMRs that survived multiple testing through bootstrapping (Figure 4) for association with schizophrenia, and also suggests a possible effect on height (Additional File 1, Figure S5).

Xu et al [18] and Srinivasan et al [19] respectively demonstrated that variants located in HARs and in regions containing NSS markers were enriched for association with schizophrenia. In our study, we compared the evolutionary enrichments of schizophrenia risk variants in DMRs, NSS markers and HARs. We validate the results of Srinivasan et al [19] (Figure 3, Figure 4). HARs do not show enrichment of disease markers by QQ plots and INRICH, unlike NSS markers and DMRs (Figure 3, Figure 4). This difference with the report of Xu et al could be due to a different freeze of the gene database used; it could also be because Xu et al. used a more stringent Hardy-Weinberg equilibrium (HWE) threshold to filter out markers from the schizophrenia GWAS [14], a step we could not replicate as the genotype data are not publicly available. We used the publicly available schizophrenia dataset that has a HWE *p*-value > 10^−6^ in controls and *p*-value > 10^−10^ in cases [14]. Interestingly, all the evolutionary annotations (DMRs, NSS markers and HARs) cover different sections of the genome with very little overlap between them (Additional File 1, Figure S2). Between the three evolutionary annotations, nearly 70,000 SNPs occur around regions with evolutionary significance (Additional File 1, Figure S2). Our results supply a wealth of information on genomic regions that are important for the evolution of humans and are also enriched for schizophrenia risk variants (NSS markers and DMRs, Additional File 3). In addition, our study provides genetic support from two independent datasets that regions which differ between modern and ancient hominids could be implicated in the development of schizophrenia. An interesting hypothesis to consider is the possibility that methylation patterns are potentially driven by the genomic sequence underneath. This hypothesis is supported by preliminary findings presented at the recently concluded World Psychiatrics Genetics Congress[41]. As such it is possible that the human specific DMRs analyzed here represent regions of the human genome where the underlying sequence might have diverged from Neanderthals and Denisovans. This hypothesis may be partially true as Gokhman et al[29] observed that some, but not all of the methylation changes were indeed driven by sequence changes. On the other hand, there also exist metastable epialleles where there are methylation differences in genetically identical individuals[42]. As such, this would suggest that not all methylation differences are driven by the underlying genomic sequence alone. We did not test whether the schizophrenia markers are human-specific or not and therefore should be investigated in future research.

Neanderthal- or Denisovan-specific DMRs showed no enrichment of association (Figure 2). This suggests that SNPs conferring vulnerability to schizophrenia occur in genomic regions whose methylation levels were altered in the modern human lineage but not in the ancestral lineages. It is possible that the evolutionary changes driving the variation in methylation status could also have made the human lineage more vulnerable to schizophrenia. A caveat to this result is that the LD structure in archaic genomes is unknown, so we cannot test LD-based enrichment in Neanderthal or Denisovan genomes. Our inter-lineage analyses with enrichment plots were thus restricted to SNPs occurring exclusively *within* DMRs. The other limitation to this comparative approach is that the GWAS data is specific to modern humans.

In previous studies [32], it was reported that 5’UTRs are the functional annotation harboring the most association with a given trait. However, the DMRs enriched for association with schizophrenia tended to localize in intronic regions (Additional File 1, Figure S4), which is in agreement with the expectation that methylation regions should not be localized in exons and UTRs. This shows that using more information to label some genomic regions in greater detail, such as potential regulatory regions in introns, might give a more precise annotation of regions of association.

Despite the genetic overlap between bipolar disorder and schizophrenia, we do not find evidence of enrichment of association of DMRs with bipolar disorder either at the SNP level (Figure 1; Additional File 1, Figure S1) or the gene level (data available on request). This could possibly be due to lack of sufficient power in the bipolar disorder GWAS [34]. Additionally, the gene-level approach utilized by INRICH depicts enrichment of association of human DMRs with height (Additional File 1, Figure S5). This evidence is lacking at the SNP level as depicted by QQ plots (Figure 1; Additional File 1, Figure S1). We speculate that this could be due to the difference in testing DMR-localized SNPs compared to genes flanking human DMRs.

Although the DMRs utilized here were obtained from bone samples, Gokhman et al [29] assert that the DMRs refer to species-specific methylation differences and not tissue-specific variations[43]. Similarly, Hernando-Herraez et al [44] noted that species-specific DMRs tend to be conserved across tissues and as such should not represent tissue-specific variations. Other studies also showed that neurological systems were enriched for methylation differences even when the tissue samples analyzed were not neurological [45–47]. Therefore, we believe that our results are valid for a ‘brain’ phenotype even though the DMRs were derived from non-brain tissues. The enrichment seen for schizophrenia also corroborates the results of Gokhman et al [29] who reported that DMRs were more enriched around genes implicated in the nervous system amongst all the organ systems tested for evolutionary changes in methylation patterns. Hernando-Herraez et al [44] also found that methylation differences between humans and great apes were located around genes controlling neurological and developmental features. It is therefore possible that the methylation differences were mediated by evolution of genomic regions controlling neurodevelopmental processes. The results of pathway analysis are consistent with this. Both the DMR and NSS regions that are enriched for association with schizophrenia contain genes that are overrepresented in ‘CREB signaling in neurons’ and ‘Synaptic long term potentiation’.

Our results hint that epigenomic evolution has taken place in genomic regions implicated in the aetiology of schizophrenia. Furthermore, these regions harbor markers that are involved in the regulation of various neurodevelopmental pathways. The fact that methylation changes also took place in these very same regions suggests a complex gene-by-environment interaction in the evolution of humans, especially for pathways that led to the development of our brain. While it is known that various factors from the environment can make long-lasting changes in DNA methylation patterns that can be subsequently inherited at a population level [30, 31, 48], the true significance of our findings from an evolutionary standpoint suggests that the superior mental abilities of our species may in part have been driven by environmental factors during the past 300,000 years [49, 50]. It is difficult to put an exact date on the emergence of the superior mental abilities that define the modern *Homo sapiens*. Anthropologists often date the onset of the advanced intellectual abilities of *Homo sapiens* from about 70,000 years onwards [51], a period which saw the emergence of art, religion [52, 53] and possibly spoken language [54]. From an evolutionary perspective, it suggests a massive leap in the animal kingdom because *Homo sapiens* became the first species not only to develop the capacity to think and imagine things that do not exist [52, 53], but also to communicate these ideas to other members of the species [54]. This ability would have been critical for effective coordination and cooperation within large groups and may even have been needed to keep a group together [55, 56]. The genomic approach to analyze mental disorders used in the present study and other studies can interrogate the effect of changes which appeared in the last 300,000 years [49, 50], but it will clearly be interesting to trace mores recent changes. If a similar method used in the reconstruction of Neanderthal and Denisovan genomes [29, 57] could be implemented on samples of ancient *Homo sapiens* from different time periods [58–65], then theoretically it should be possible to reconstruct the methylomes and regions of recent evolution from ancient humans [66]. Subsequently, more detailed ‘time-course’ analyses of changes in methylation patterns and in other regions of recent evolution and their implications in schizophrenia will surely result in more detailed elucidation of the evolutionary hypothesis of schizophrenia. This is the promise of the novel field of paleoepigenetics that seeks to infer past environmental cues that affected the epigenomes of ancient individuals[42, 43].

## Conclusions

In summary, we have demonstrated that human genomic regions whose methylation status was altered during evolution are enriched in markers that show association with schizophrenia. Our results concur with previous genomic studies demonstrating that methylation changes in *Homo sapiens* have had the greatest impact on the nervous system. and provide evidence that epigenomic evolution plays a role in conferring a high risk of schizophrenia on humans. Future research should attempt to perform a finer temporal resolution of the origins of psychosis through the prism of evolutionary epigenomics. To explore the period of evolution before the *Homo* lineage, it would also be interesting to determine whether methylation signatures from primates are enriched for schizophrenia markers. Future research should also investigate the influence of human-specific DMRs on height.

## Methods

### Differentially Methylated Region data

Coordinates for DMRs were obtained from data publicly available in Supplementary Table S2 of Gokhman et al, 2014 [29]. This file contained DMRs inferred by comparing genome sequence of fossilized Neanderthal and Denisovan limb samples with methylation data from osteoblasts of modern humans. From the genomes of the Neanderthal and Denisovan samples, Gokhman et al inferred methylation by utilizing the natural degradation of methylated cytosine (C) to thymine (T) to create a C→T ratio [29]. The methylation information, in the form of C→T ratio, was then compared with each of the three species and classified according to the hominid in which the methylation change occurred, i.e. human-specific, Neanderthal-specific and Denisovan-specific DMRs. These DMRs do not represent tissue-specific methylation but species-specific methylation [29]. The human-specific DMRs comprise regions that have both gained and lost methylation in comparison to Neanderthal- and Denisovan-specific DMRs. DMRs that could not be classified reliably in any of the three species (unclassified DMRs) [29] were not used. Full methodological details for assigning DMRs are in the Supplementary File of the original paper [29].

### HAR data

Genomic coordinates were obtained from publicly available data (docpollard.com/2x) for three classes of human accelerated region: HARs, in which regions conserved in mammals are accelerated in humans; PARs, in which regions conserved in mammals are accelerated in primates; and pHARs, in which regions conserved in primates (but not other mammals) are accelerated in humans. Conversion to hg19 assembly was performed using the liftOver tool from the UCSC Genome Browser.

### NSS data

NSS data was obtained as a list of markers with corresponding NSS values from Srinivasan et al [19]. Markers with negative values, indicating positive selection in humans, were filtered out and used for analysis.

### GWAS data

Summary statistics from GWAS of 12 common traits were obtained from published datasets: schizophrenia (SCZ) [14], bipolar disorder (BPD) [34], attention deficit hyperactivity disorder (ADHD) [35], rheumatoid arthritis (RA) [36], blood lipid markers (high density lipoprotein (HDL), low density lipoprotein (LDL), triglycerides (TG), total cholesterol (TC)) [37], blood pressure (systolic blood pressure (SBP), diastolic blood pressure (DBP)) [38], body mass index (BMI) [39], and height [40]. For studies published with hg18 coordinates (BPD, SBP, DBP, HDL, LDL, TG, TC, ADHD, RA), conversion to hg19 was performed using the command line version of the liftOver tool from the UCSC Genome Browser (http://hgdownload.cse.ucsc.edu/downloads.html#utilities_downloads). For BMI and height SNPs, the genomic coordinates were obtained by mapping them to the assembly of 1000 Genomes Project Phase 1 reference panel SNPs [67].

### SNP assignment to DMRs

SNPs were assigned to DMRs with LDsnpR [68] using positional binning and LD (linkage disequilibrium)-based binning in R [69]. We used both methods because DMR-localized SNPs that were not genotyped in a specific GWAS would be missed if we used positional binning alone [68] (Additional File 1, Table S1). The LD file utilized in HDF5 format was constructed on the European reference population of 1000 Genomes Project and can be publicly downloaded at: http://services.cbu.uib.no/software/ldsnpr/Download.

### Enrichment analyses with stratified Quantile-Quantile (QQ) Plots

QQ plots are an effective tool to visualize the spread of data and any deviations from the expected null distributions. They are frequently utilized in GWAS to depict enrichment of true signals. When the observed distribution of data matches the expected distribution, there is a lack of enrichment and a line of equality is obtained that depicts the null hypothesis. A distribution such as this reflects no enrichment of observed over expected data distribution. However, if the observed and expected distributions differ, there will be deviation from this null line. As described in detail by Schork et al [32], leftwards deflections from this null line represent enrichment. The higher the leftward deflection, the greater is the enrichment of true signals. In GWASs, due to the extremely low *p*-values of SNPs, it is common to depict *p*-values by converting them to negative log_10_ values so that smaller *p*-values give higher negative logarithmic values. We plotted the negative log_10_ of the observed *p*-values of SNPs against the expected negative log_10_ of a normal distribution. The distributions were corrected for genomic inflation by *λ*_GC_. This method of enrichment was used to show for example [32] that specific genomic regions are enriched for trait-associated SNPs and are much more likely to associate with a given trait than SNPs distributed across a genome. In other words, when SNPs are stratified according to specific genomic regions, there is a greater enrichment of true signals than what is observed in the GWAS. Using a similar approach, we binned SNPs that fall in DMR regions and plotted the stratified *p*-value distribution.

### Enrichment analyses with INRICH

The stratified QQ plots are a useful visual tool for observing the presence or absence of enrichment of true signals in a given set of SNPs. However, to quantify the enrichment visually observed, we used the INterval EnRICHment Analysis (INRICH) tool. It is a robust bioinformatics pipeline to determine enrichment of genomic intervals implicated by LD with predefined or custom gene sets [33]. It takes into account several potential biases that can otherwise lead to false positives. It is well suited for testing GWAS-implicated SNPs for association with gene sets as it controls for variable gene size, SNP density, LD within and between genes, and overlapping genes with similar annotations. We followed the procedure described by Xu et al [18], with the extended MHC region (chr6:25-35Mb) masked and SNPs with minor allele frequency (MAF) <0.05 excluded. Full details may be found in Additional File 1.

### Pathway analysis

Pathway analysis was performed using Ingenuity Pathway Analysis (IPA) from QIAGEN (www.qiagen.com/ingenuity, last accessed 26^th^ August 2016). The reference set was Ingenuity Knowledge Base (Genes). Both direct and indirect relationships were analyzed. All data sources were included with the confidence parameter set to experimentally observed and highly predicted pathways for Human. Under ‘Tissues & Cell Lines’, we performed the analysis once with all organ systems and once with only the nervous system. 5338 enriched DMR SNPs in 329 enriched DMRs (Additional File 3) were mapped to 349 unique RefSeq genes and 446 RefSeq genes in LD using the method of Schork et al [32]. Genes in LD blocks containing enriched NSS markers were determined in a similar manner. 4276 enriched NSS markers mapped to 648 overlapping RefSeq genes and 1363 RefSeq genes in LD. IPA was performed on these gene lists.

## Additional Files

Additional File 1: Additional Method, Figures and Tables

Additional File 2: Annotation of all DMRs with schizophrenia-associated SNPs. This file contains annotation of all the human-lineage specific DMRs that are associated with schizophrenia markers. Details of the various markers present within each DMR is provided, along with the marker with most significant *p*-value.

Additional File 3: Annotation of enriched DMRs with genes, promoters, CpG islands and enhancers. This file contains detailed annotation of those human-lineage specific DMRs that are enriched for association with schizophrenia markers(except those in the MHC region). Compared to Additional File 2, these DMRs represent those that are enriched and whether they are present in any genes, promoters, enhancers or CpG islands.

## Abbreviations

(3’UTR): 3’ untranslated region,
(5’UTR): 5’ untranslated region,
(ADHD): attention deficit hyperactivity disorder,
(BMI): body mass index,
(BPD): bipolar disorder,
(CpG): 5’ Cytosine-phosphate-Guanine 3’,
(CREB): cyclic adenosine monophosphate responsive element binding protein,
(DBP): diastolic blood pressure,
(DMR): differentially methylated region,
(DNA): deoxyribonucleic acid,
(GWAS): genome-wide association studies,
(HARs): Human Accelerated Regions,
(HDL): high density lipoprotein,
(HWE): Hardy-Weinberg equilibrium,
(INRICH): Interval enRICHment analysis tool,
(IPA): Ingenuity Pathway Analysis,
(LD): linkage disequilibrium,
(LDL): low density lipoprotein,
(MAF): minor allele frequency,
(MHC): Major histocompatibility complex,
(NSS): Neanderthal Selective Sweep,
(QQ): quantile-quantile,
(RA): rheumatoid arthritis,
(SBP): systolic blood pressure,
(SCZ): schizophrenia,
(SNP): single nucleotide polymorphism,
(TC): total cholesterol,
(TG): triglycerides.

## Declarations

### Ethics Approval and consent to participate

Not applicable

### Consent for publication

Not applicable

### Availability of Data and Materials

The code supporting the results of this article is available in the Zenodo repository at http://doi.org/10.5281/zenodo.198451. GWAS datasets, DMR data and HAR data are publicly available as described in the Methods section.

### Authors’ contributions

NB carried out the bioinformatics analyses, contributed to the design of the study and drafted the manuscript. TP, SG, FB contributed to statistical and LD analyses. OAA and VMS critically revised the manuscript. SLH conceived of the study, participated in its design and coordination, and helped to draft the manuscript. All authors read and approved the final manuscript.

## Acknowledgements

We thank Profs. Anders Dale and Wesley Thompson, University of California, San Diego for helpful discussions and Isabel Hanson Scientific Writing for formatting and submission. The code used to generate the QQ plots was graciously made publicly available by Matthew Flickinger PhD, University of Michigan at http://genome.sph.umich.edu/wiki/Code_Sample:_Generating_QQ_Plots_in_R. We thank Ke Xu from Icahn School of Medicine, Mount Sinai for providing the URL for downloading the HAR datasets. We thank the author of the book *Sapiens: A Brief History of Humankind*, Yuval Noah Harari, PhD, for some of the ideas discussed in the paper. Finally, we thank the reviewers for their time and their suggestions for improving the manuscript.

